# Phylogenetic analysis of migration, differentiation, and class switching in B cells

**DOI:** 10.1101/2020.05.30.124446

**Authors:** Kenneth B. Hoehn, Oliver G. Pybus, Steven H. Kleinstein

## Abstract

B cells undergo rapid mutation and selection for antibody binding affinity when producing antibodies capable of neutralizing pathogens. This evolutionary process can be intermixed with migration between tissues, differentiation between cellular subsets, and switching between functional isotypes. B cell receptor (BCR) sequence data has the potential to elucidate important information about these processes. However, there is currently no robust, generalizable framework for making such inferences from BCR sequence data. To address this, we develop three parsimony-based summary statistics to characterize migration, differentiation, and isotype switching along B cell phylogenetic trees. We use simulations to demonstrate the effectiveness of this approach. We then use this framework to infer patterns of cellular differentiation and isotype switching from high throughput BCR sequence datasets obtained from patients in a study of HIV infection and a study of food allergy. These methods are implemented in the R package *dowser*, available at https://bitbucket.org/kleinstein/dowser.

**Author summary:** B cells produce high affinity antibodies through an evolutionary process of mutation and selection during adaptive immune responses. Migration between tissues, differentiation to cellular subtypes, and switching between different antibody isotypes can be important factors in shaping the role B cells play in response to infection, autoimmune disease, and allergies. B cell receptor (BCR) sequence data has the potential to elucidate important information about these processes. However, there is currently no robust, generalizable framework for making such inferences from BCR sequence data. Here, we develop three parsimony-based summary statistics to characterize migration, differentiation, and isotype switching along B cell phylogenetic trees. Using simulations, we confirm the effectiveness of our approach, as well as identify some caveats. We further use these summary statistics to investigate patterns of cellular differentiation in three HIV patients, and patterns of isotype switching in an individual with food allergies. Our methods are released in the R package *dowser*: https://bitbucket.org/kleinstein/dowser.

## Introduction

The adaptive immune system in humans depends on B cells to produce antibodies capable of neutralizing a wide array of pathogens. Antibody structures are initially expressed as B cell receptors (BCRs) on the surfaces of B cells. BCRs are generated through random V(D)J recombination and then subjected to repeated rounds of somatic hypermutation (SHM), cell proliferation, and selection for antigen binding [1]. This evolutionary process, called affinity maturation, creates many lineages of B cells that each descend from a single naïve progenitor cell. Cells within a clonal lineage differ predominately by point mutations. The genetic variation within these clonal lineages has been long investigated using phylogenetic methods [2]. When obtained through high throughput sequencing, BCR sequences have shown promise in elucidating information about the adaptive immune response in humans, such as the sequence of mutations that occur during antibody co-evolution with HIV [3], and the process of mutation and selection during affinity maturation generally [4]. Other important biological processes may occur as BCR sequences evolve, such as B-cell migration between tissues [5], differentiation into cellular subsets [6], and antibody isotype class switching [7]. If these processes co-occur with SHM, then in principle they can be investigated and inferred using phylogenetic techniques.

Migration and cellular differentiation in B cells can be viewed as analogous to geographic spread of rapidly evolving viruses, the study of which – viral phylogeography – has advanced both in theory and application in the past decade (e.g. Lemey *et al*. 2009). For example, phylogeographic methods have been used to determine the origin of the HIV pandemic [9], factors influencing the recent Ebola epidemic [10,11], and the epidemic spread of Zika virus [12,13]. Modern phylogeographic analyses typically model phylogenetic sequence evolution [14], and changes in geographic location within a unified framework [15,16]. Successfully developing a phylogeographic framework for B cell lineages would enable the testing of new hypotheses regarding the nature of evolution during affinity maturation.

There are serious challenges that must be addressed before using modern phylogeographic methods on B cell repertoire datasets. Such techniques typically rely on molecular clock trees, whose branch lengths represent elapsed time between nodes [15]. Accurately modelling sequence change through time requires either data sampled at multiple time points, or prior information about expected rate of sequence evolution. These are not frequently available for B cell lineages. Data samples, particularly biopsies, are often only taken at a single time point [5,17], and the variation of B cell mutation rate over time is largely unknown and likely dependent on cell subset. Even using a Markov model to describe state changes along B cell molecular phylogenies is not straightforward: B cell lineage trees frequently contain identical sequences with different states. This results in state changes across zero-length branches, which are not able to be fit within a Markov model framework. Further, modern phylogeographic techniques often rely on Markov chain Monte Carlo sampling, which makes them computationally intensive and impractical to apply to thousands of sequences. Unfortunately, B cell bulk repertoires often contain millions of sequences, and individual lineages sometimes contain thousands of unique sequences.

We propose that hypotheses about B cell migration and differentiation may be usefully investigated using heuristic summary statistics that characterize the distribution of trait values along phylogenetic trees. Indeed, such heuristic approaches which do not depend on branch length have historically been a popular means of testing hypotheses about migration between populations [18–20]. While tree based summary statistics have been previously used to assess B cell migration [5], differentiation [6], and isotype switching [7], these approaches have not been tested through simulations and their general accuracy is unclear. To address this methodological gap, we develop a set of maximum parsimony-based statistics that summarize the relative distribution of B cell states along lineage trees within repertoires and introduce a framework for assessing the significance of their difference from randomized trees. We demonstrate through simulations that these tests relate intuitively to different regimes of migration and differentiation. To demonstrate its utility, we use this framework to test hypotheses regarding differentiation of cell types in HIV infection, and sequential class switching to IgE and IgG4. We introduce a statistically principled and scalable means of analyzing the evolution of discrete traits in B cell repertoires. We release these methods in the R package *dowser*.

## Methods

### Predicting states of internal tree nodes

The goal of the discrete trait analysis framework presented here is to characterize the distribution of predicted trait values along B cell lineage trees. Given an alignment of sequences inferred to descend from the same naïve ancestor (i.e. the same clonal family), lineage tree topologies and branch lengths were estimated using maximum parsimony using *dnapars* v3.967 [21]. Importantly, the statistics presented here are not limited to tree topologies inferred through maximum parsimony.

Maximum parsimony is also used to infer the discrete character states (e.g. cell subtype, isotype, tissue) of internal nodes, given a tree in which each tip is associated with a given character state. Nodes with different states from their immediate ancestors are counted as state changes. More specifically, internal node states were reconstructed using the Sankoff dynamic programming maximum parsimony algorithm [22], which, given a weight matrix for each type of state change, determines the minimum number of state changes that must be made along the tree given the states at the tips. The backtrace step of this algorithm can be used to determine a set of most parsimonious internal node states. Often there are multiple such maximum parsimony sets. To represent state changes across ambiguous internal node sets, trajectories with equal parsimony were randomly chosen in the backtrace step of the Sankoff algorithm, beginning at the root of the tree and moving towards the tips. This process is performed 100 times for each tree, and the mean of each type of state change was reported.

Strictly bifurcating B cell lineage trees frequently have clusters of nodes separated by zero-length branches (soft polytomies), which represent a high degree of uncertainty in tree topology. This uncertainty in the order of bifurcating nodes can result in a potentially large number of uninformative state changes along the polytomy. Multiple steps were taken to minimize the effects of random polytomy resolution (**Supplemental File S1**). Briefly, nodes within each polytomy were first re-ordered to minimize the number of state changes along the tree. To represent the uncertainty in the order of state changes, nodes within each polytomy were grouped together into separate subtrees according to their predicted state. These state-specific subtrees were then joined together in a balanced manner, ensuring that state changes could occur in any direction among the states contained within the polytomy (**Supplemental File S1**).

### Testing trait-phylogeny association

Analysis begins with a B-cell lineage tree topology with discrete character states (trait values) associated with each tip, and internal node states reconstructed through maximum parsimony. The goal of our discrete trait analysis framework is to determine how the distribution of discrete character states along the internal nodes of a tree differs from its expectation if traits are randomly distributed among the tips. The statistics introduced herein are shown graphically in **Fig 1**. More formally, if there are *m* possible discrete character states, and *o*_*ij*_ is the number of state changes from type *i* to type *j*, the three statistics investigated are defined as:

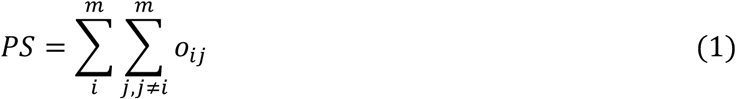

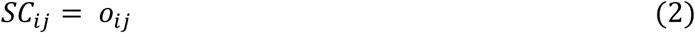

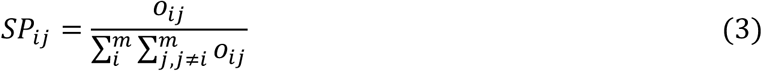

**Fig 1:**
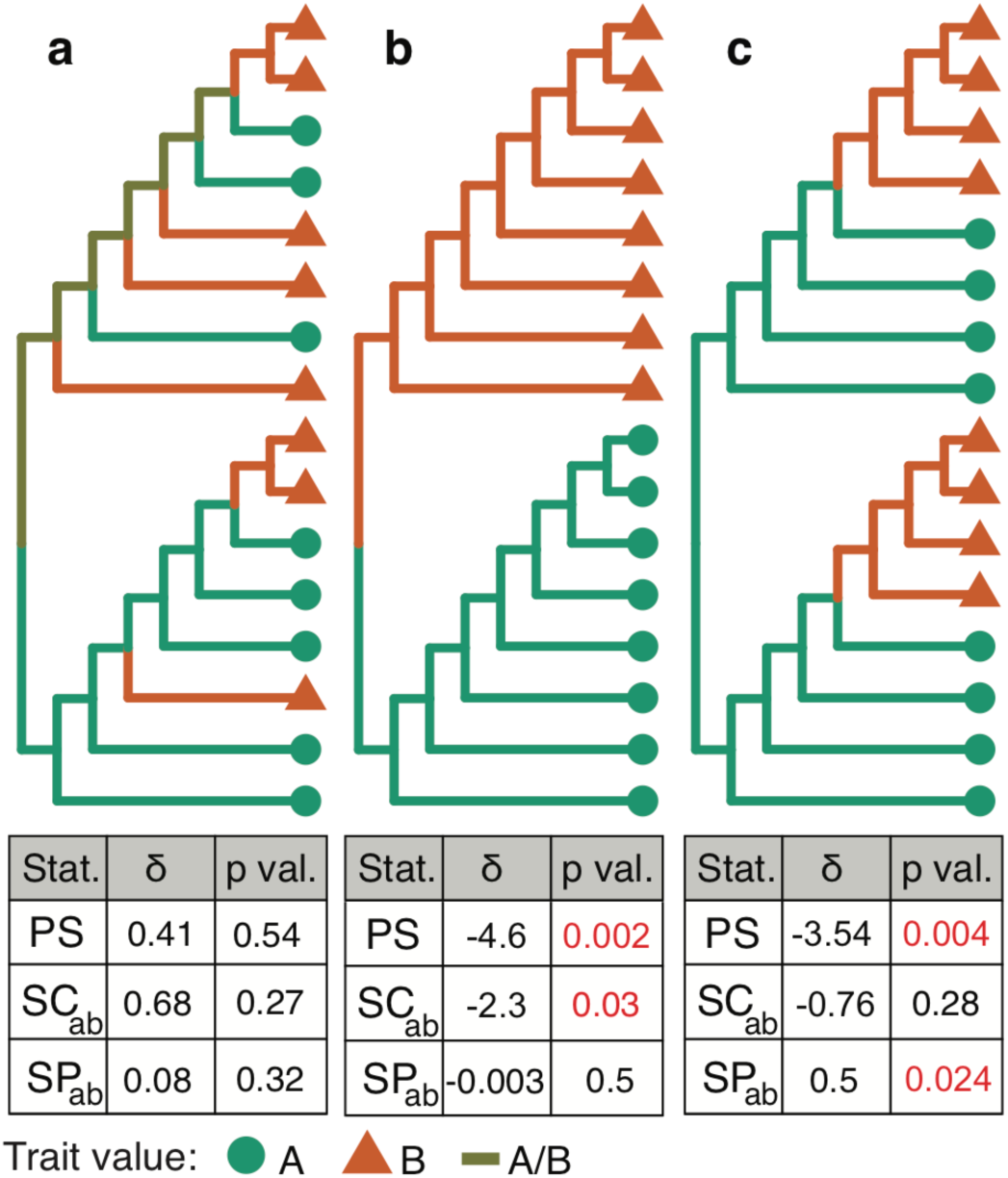
Hypothetical phylogenies used to illustrate tree/trait association statistics. Trait values at internal nodes of the tree are predicted using maximum parsimony reconstruction given the trait values at the tips, which are shown using different colors and shapes. Below each tree is a table of **δ** and corresponding *p* values for *PS, SC*, and *SP* tests performed on each tree, calculated using 1000 permutations. Tests were performed on the tree topologies themselves – bootstrap replicates were not performed. **(a)** *No association between tip-trait values and tree*: Distribution of traits across this tree is indistinguishable from randomly distributed traits by any statistic used. **(b)** *Tip-trait values clustered in tree*: Association between trait and tree structure revealed by significantly low *PS* statistic. This tree also has a significantly low switch count statistic from *A* to *B* (*SC*_*ab*_). Further, this tree has an identical switch proportion statistic (1/2) from *A* to *B* as from *B* to *A*, which is not significantly different from permuted data (*SP*_*ab*_). **(c)** *Biased ancestor/descendant relationships among trait values*: As in **b**, this tree also shows a significant relationship between tree and trait distribution (*PS*). However, this tree also has a significantly high *SP* statistic from *A* to *B* (*SP*_*ab*_). Only the *SP* test captures the directionality of this relationship.

The *PS* (parsimony score) statistic is the total number of state changes along a tree. The *SC* (switch count) statistic from *i* to *j* is the number of state changes from state *i* to *j*. The *SP* (switch proportion) statistic from *i* to *j* is the proportion of state changes from state *i* to *j*.

We calculate the significance of these three statistics using a permutation test. This is done by randomizing traits at the tips of the lineage tree, re-calculating each statistic on the permuted tree, and repeating for a specified number of replicates. For each replicate, we calculate δ, which is the difference between the statistic calculated on the observed tree and the same statistic calculated on the permuted tree. If mean δ > 0 (hereafter mean δ is indicated by **δ**), this indicates the statistic is on average higher in observed trees than in permuted trees. For a one-tailed test, we calculate the *p* value that **δ** > 0 as the proportion of replicates in which δ ≤ 0. Similarly, we calculate the *p* value that **δ** < 0 as the proportion of replicates in which δ ≥ 0. For a two-tailed test, we calculate the *p* value that **δ** > 0 as the proportion of replicates in which δ < 0, plus half of the replicates in which δ = 0. We refer to the calculation of these *p* values for the statistics in **Eq. 1-3** as the *PS* test, *SC* test, and *SP* test, respectively.

These three statistical tests capture different aspects of how the distribution of characters observed along a lineage tree differs from random association between tree topology and trait values. The *PS* test determines the extent to which trait values are clustered together within the tree. A significantly low *PS* statistic (i.e. **δ** < 0, *p* < 0.05) indicates identical trait values are more closely clustered together within the tree than expected from random association between tree topology and trait values. By contrast, a significantly high *PS* statistic indicates identical trait values are less clustered together than expected by chance. Variations of this test were previously developed in [18] and applied in [23] to study spread of influenza.

While the *PS* test only determines a general association between trait values and tree topology, the *SC* and *SP* tests are both aimed at determining whether a particular trait value is more ancestral to another in the tree. A significantly high *SC* statistic (**Eq. 2**) from state *i* to state *j* indicates a greater number of switches from state *i* to state *j* than expected from random association between tree topology and trait values. The *SC* test was used by [19] in the context of virus phylogeography, and by [6] to decompose the phylogenetic relationships among B cell subtypes within HIV infection. The *SC* test, however, is not purely a metric of whether state *i* tends to be more immediately ancestral to state *j* than expected. This is because trees with randomized tip states often have more state changes events in general, hence **δ** values tend to be negative even when there is no polarity in the ancestor/descendant relationship among type *i* and *j* (**Fig 1B**).

To normalize for changes in the total number of state changes between observed and permuted trees, we introduce the *SP* test. A significantly high *SP* statistic (**Eq. 2**) from state *i* to state *j* indicates a greater proportion of switches from state *i* to state *j* than expected from a random distribution of trait values at the tips. In contrast to the *SC* test, a significant association between the tree and trait may exist, but only associations in which one trait is more often ancestral to the other will give rise to a significantly high or low *SP* statistic (**Fig 1C**). Further, the denominator of the *SP* statistic can be altered to test other hypotheses. For instance, to test whether a greater proportion of state changes to state *j* come immediately from state *i* than expected by chance, one can restrict the analysis to consider only state changes towards *j*.

### Accounting for uncertainty in tree topology

To account for uncertainty in tree topology, we bootstrap multiple sequence alignments within each clone [24]. This is performed by random sampling with replacement of the columns of a multiple sequence alignment. Lineage tree topology and branch length estimation then proceeds as before. Test statistics are calculated for each bootstrap replicate tree, and then for a single permutation of the traits at that tree’s tips. Calculations for **δ** values proceed as before, but with each *o*_*ij*_ value indexed for each replicate. This procedure is very similar to that proposed in [25] and can in principle be extended to other sets of tree topologies, such the posterior distribution of tree topologies generated by MCMC sampling under Bayesian phylogenetic inference.

### From trees to repertoires

B cell repertoire datasets often consist of hundreds or thousands of B cell lineage trees. Often hypotheses do not concern individual lineages, but instead the behavior of the collection of B cell lineages as a group. To characterize multiple B cell lineages, the observed and permuted summary statistics are summed across all lineages for each bootstrap replicate. Additionally, traits may be permuted among trees, which may increase statistical power and detect nonrandom association among trait types within trees.

### Simulations

We tested the performance of the three proposed statistics using simulations based on B cell lineage trees estimated from an empirical dataset. Sequences were obtained from peripheral blood samples taken from one subject at ten time points, from eight days before to 28 days after influenza vaccination [26] (subject 420IV). Sequence preprocessing and clonal clustering are described in [4]. Sequences were down-sampled by 50%, and only clones with >10 unique sequences were retained. A total of 399 clones containing 11 to 370 (mean=26.2) unique sequences remained. Tree topologies and branch lengths were estimated for each clone using *dnapars* v3.967 [21] via the R package Alakazam v0.3.0 [27]. We simulated state changing down each tree using a Markov model parametrized by initial frequencies ***π*** for each state, relative rate parameters *r*_*ij*_ for each pair of possible states *i* and *j*, and *r*, the average rate of state changes per mutation per site. The mean value of this rate matrix was calculated as the sum of the diagonal elements weighted by their initial state frequencies. All values of the matrix were then divided by this mean and multiplied by *r*. This calibration was performed so that *r*\**l* state change events are expected to occur across a branch of length *l* mutations/site. For each tree, the state at the germline node was randomly drawn based on each state’s *π* value. For each node after the germline, the rate matrix is multiplied by the node’s ancestral branch length and exponentiated to give the probability of each state at the descendant node, given the state at the node’s immediate ancestor. The state at the descendant node is randomly chosen based on these probabilities. This process begins at the germline node and continues down the tree until each tip node has a state. Because each tip corresponds to a sequence, this forms a dataset of sequences paired with simulated discrete characters. Internal node states are not included in the final simulated dataset used for analysis. Simulations were performed with two state models (*A* and *B*) that explored a large parameter space (*π*_*a*_ = 0.5, 1; *r*_*ab*_ = 0.1, 1, 10; *r* = 10, 25, 50, 100, 1000), and four state models (*A, B, C, D*) that explored more complex patterns of state change at low overall rate (*r* = 10). Twenty simulation repetitions were performed for each parameter combination. Statistical tests were performed as described in **Methods**; however, to improve computational efficiency simulation analyses did not use bootstrapped multiple sequence alignments, and instead performed 100 permutations on a fixed maximum parsimony tree for each clone. Only clones with more than one state type were analyzed.

### Empirical datasets

We demonstrate the utility of the proposed discrete trait framework by analyzing two empirical datasets. The first was aimed and understanding B cell differentiation during HIV infection, and consists of BCR mRNA sequences taken from sorted populations of unswitched memory B cell (MBC), CD19^hi^ MBC, CD19^lo^ MBC, and germinal center B cells (GCBC) from three HIV viremic subjects (subject 1-3; [6]) Each dataset was subsampled to a maximum of 50,000 total sequences, and only clones with more than 10 sequences were retained. Unique sequences associated with more than one cell type were kept distinct. This resulted in 128, 197, and 174 clones with a mean of 53, 38.6, and 31.8 unique sequences per clone, for subjects 1-3 respectively. State changes across all lineages for each subject were calculated over 100 bootstrap replicates.

The second dataset was aimed at understanding isotype switching patterns in human children, and consists of BCR mRNA sequences obtained from peripheral blood samples taken from a human child each year from age 1 to 3 years old [28]. Preprocessing, including grouping of sequences into clonal clusters, is detailed in **Supplemental File S2**. Only clones with at least 4 unique sequences and more than one isotype were retained. Unique sequences associated with more than one isotype were kept distinct so each sequence was associated with one isotype. This resulted in 768 clones with a mean of 9.3 unique sequences each. State changes across all lineages were calculated over 100 bootstrap replicates.

## Results

We outline three parsimony-based summary statistics to characterize the distribution of trait values along B cell lineage trees (**Fig 1**). The significance of these statistics can be tested by comparing observed values within the set of trees that comprise a repertoire to those obtained from permuting trait values at the tree’s tips. The first statistic, the parsimony score (*PS*), is the total number of trait value state changes that occurred along a lineage tree. A *PS* test with **δ** < 0 and *p* < 0.05 (*i*.*e*. a significantly low *PS* statistic) indicates the trait values cluster together in the observed trees more often than expected by chance (**Fig 1**). We propose two other statistics aimed at determining whether one state is more frequently the immediate ancestor to another state than expected by chance. The switch count (*SC*) from state *i* to *j* is the number of state changes that occurred from *i* to *j* [19], while the switch proportion (*SP*) from state *i* to *j* is the proportion of state changes that occurred from *i* to *j*. An *SC* or *SP* test from *i* to *j* with **δ** > 0 and *p* < 0.05 indicates trait value *i* was more frequently immediately ancestral to state *j* than expected by chance. We expect the *SP* test to be more sensitive to this relationship than the *SC* test because it accounts for the increased number of state changes expected in randomized trees (**Fig 1B-C**). Similarly, an *SP* test from *i* to *j* with **δ** < 0 and *p* < 0.05 indicates trait value *i* was less ancestral to state *j* than expected (**Fig 1B**). All three of these tests may be used to characterize individual lineages or entire B cell repertoires; in this paper we will focus exclusively on repertoires.

### Differentiating state change patterns with two states

We used simulations to test the performance of our proposed tests. We model B cell migration/differentiation using a Markov model with two states, *A* and *B*, and empirically-derived linage tree topologies (**Methods**). Briefly, the pattern of state changes along a tree was determined by the probability that the state at the root was *A* (*π*_*a*_ = 0.5, 1; *π*_*b*_ *= 1 – π*_*a*_), the average rate of state change (*r* = 10, 25, 50, 100, 1000 changes/mutation/site), and the relative rate of change from *A* to *B* (*r*_*ab*_ = 0.1, 1, 10; *r*_*ba*_ *= 1/r*_*ab*_). These parameters represent a range of slow, fast, biased, and unbiased state change patterns along a B cell lineage. Each simulation resulted in a dataset of BCR sequences, each associated with a single trait value (*A* or *B*) resulting from the simulation process. The goal of our simulation analysis is to determine if the summary statistics provide useful information about the mode and tempo of trait evolution.

We ran 20 simulation repetitions for each parameter combination, and tested the significance of each of the proposed statistics to assess their statistical power. Our simulations are designed to generate trees whose tip-states are more clustered together than if the tips states are randomly distributed across the tree tips. Consistent with this expectation, 320/320 simulation repetitions in which *r* < 1000 (*i*.*e*. overall rate of state change < 1000 changes/mutation/site) showed a significantly low *PS* statistic regardless of other parameters (**δ** < 0; one-tailed *p* < 0.05; **Supplemental File S3**). This confirms the *PS* test’s usability for detecting nonrandom association between tree topology and trait values. However, at *r* = 1000, only 3/80 repetitions showed a significantly low *PS* statistic (**Supplemental File S3**), indicating this relationship is difficult to detect at high rates of state change.

We used the same simulations to test whether the *SC* statistic was capable of detecting the direction of state changes in B cell repertoires. A total of 300 simulation repetitions were performed using parameters expected to give biased (directed) state changes; namely, with lineages always beginning in *A* (*π*_*a*_ = 1) and/or highly biased rates of state change from *A* to *B* (*r*_*ab*_ = 10). Surprisingly, only 3/300 of these simulations showed a significantly high *SC* from *A* to *B* (**δ** > 0; one-tailed *p* < 0.05; **Supplemental File S4**). By contrast, 186/300 showed a significantly low *SC* from *A* to *B* (**δ** < 0; *p* < 0.05). This indicates that significantly high *SC* statistics are highly conservative, while significantly low *SC* statistics are primarily driven by overall phylogenetic association with a trait. This issue is likely exacerbated as dataset size grows, hence the *SC* test is likely still useful for single lineages [19,20] or for detecting very strong trends in large datasets [6]. However, given these results the *SC* test does not appear appropriate as a general solution for detecting biased migration and differentiation in B cell repertoire datasets.

We next tested whether biased state change patterns were detected by the *SP* test. To test this method’s false positive error rate, we first investigated simulations with totally unbiased state changes; namely, in which lineage trees were equally likely to begin at state *A* as *B* (*π*_*a*_ = 0.5) and had equal rates of state changes between *A* and *B* (*r*_*ab*_ = *r*_*ba*_ = 1). *SP* tests from *A* to *B* on these datasets resulted in a roughly uniform distribution of *p* values at all tested migration rates (**δ** > 0, *p* < 0.05 in 5/100; **Fig 2A**). This indicates that completely unbiased state changing is consistent with the null hypothesis of this test. Simulations in which lineages always had state *A* at the root (*π*_*a*_ = 1) and/or the relative rate of state change was higher from *A* to *B* (*r*_*ab*_ = 10) were expected to give high *SP* statistics. At low overall rates of state change (*r* = 10), 55/60 of these simulations had significantly high *SP* statistics from *A* to *B* (**δ** > 0; *p* < 0.05; **Fig 2B-D**). At higher rates of state change (*r* = 25, 50, 100, or 1000), this relationship diminished in these simulations as the distribution of trait values became less distinguishable from random association (**Fig 2B-D)**. These results indicate that, under this two state Markov model framework, a significantly high *SP* statistic is associated with biased origination, biased rate of state change, or both depending on the overall rate.

**Fig 2:**
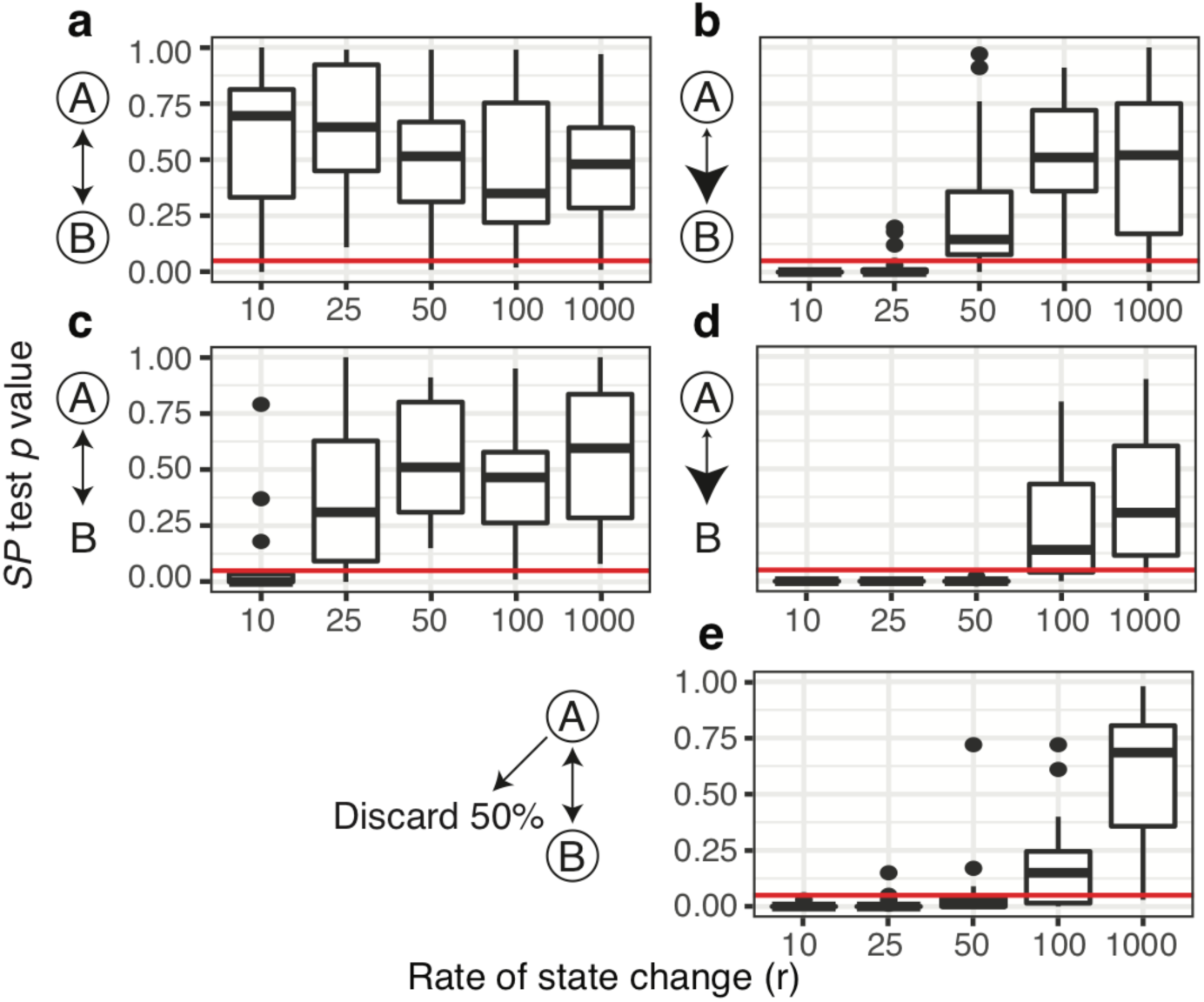
Distribution of *SP* test *p* values from *A* to *B* from two state simulation analyses in which state change between state *A* and *B* was determined by the probability of starting in *A* (*π*_*a*_), relative rate of migrating from *A* to *B* (*r*_*ab*_), and the average rate of state change (*r*). To the right of each plot, possible starting states are circled, relative rates are shown by arrowhead size. **(a)** *π*_*a*_ = 0.5, *r*_*ab*_ = 1, fully unbiased state change, shows roughly uniform distribution of *p* values at all tested rates. (b) *π*_*a*_ = 1, *r*_*ab*_ = 1 shows low *p* values at low rates (10) but not at higher rates. **(c)** *π*_*a*_ = 0.5, *r*_*ab*_ = 10 shows low *p* values at rates < 50. **(d)** *π*_*a*_ = 1, and *r*_*ab*_ = 10 shows low *p* values at rates < 100. **(e)** *π*_*A*_ = 1, *r*_*ab*_ = 1 shows low *p* values at rate < 50 if 50% of *A* sequences are discarded. Compared to **(a)**, this shows that *p* values are sensitive to biased sampling of sequences. Red lines show the cutoff of *p* value = 0.05.

Finally, we used these simulated datasets to test whether the *SP* test is affected by biased data sampling, as this potential bias is important for some other phylogeographic methods of trait evolution e.g. [29]. We tested this by randomly discarding half of the sequences associated with *A* in simulations with totally unbiased state change (*π*_*a*_ *=* 0.5, *r*_*ab*_ = *r*_*ba*_ = 1). Though *SP* tests from *A* to *B* on these datasets gave a uniform *p* value distribution when all sequences were included (**Fig 2A**), *SP* statistics became significantly high when half of *A* sequences were discarded (**Fig 2e**). This indicates that severely biased sampling may give a similar signature as biased origination or state change for the *SP* test (**Figs 2B-D**). Biased sampling may be caused by a variety of experimental factors, and applications of these statistics to empirical datasets will need to carefully consider possible effects of biased data collection for each trait type.

### Differentiating complex relationships among trait values

All the tests detailed above are extendable to data with more than two states; however, due to its superior performance in two state simulations, we will focus in the rest of this study on the *SP* test. The permutation step of the *SP* test usually permutes trait values within each tree separately (**Methods**). However, when more than two states are present it may be advantageous to randomize trait value assignments among trees rather than just within each tree. This changes the null hypothesis, which is now that the proportion of state changes observed is the same as that expected if trait values are randomly distributed among all trees. Deviations from this null hypothesis may be due not only to biased ancestor/descendant relationships within individual trees, but also co-occurrence of trait values within different trees. To demonstrate the difference between these two mechanisms, we performed simulations with four trait values: *A, B, C*, and *D*. To test the difference between simple association and biased ancestry, these simulations used unbiased state change between *A* and *B*, and unidirectional state change from *C* to *D*. Trees began with states *A, B*, or *C* in equal probability; state changes were allowed in both directions from *A* to *B* and unidirectionally from *C* to *D*. For each repetition, the rate of state change (*r*) = 10, and relative rates were equal among allowed state changes. Performing the *SP* test on these simulations while permuting among trees showed significantly higher *SP* statistics in both directions between *A* and *B*, and between *C* and *D* than expected (20/20 for each; **Fig 3A**). This indicates the *SP* test when permuting among trees detected the association between these trait values but not the directionality of *C* to *D* state changes. In contrast, the *SP* test when permuting only within trees correctly yielded a significantly high *SP* statistic from *C* to *D* in 19/20 simulations; further, no simulation yielded a significantly high *SP* statistic from *D* to *C*, indicating a low false positive rate. No simulation using either permutation method showed a significantly high *SP* statistic between unassociated trait values (e.g. *A* and *C*), indicating a low false positive rate. These results indicate that permuting trait values within trees is a more effective means of detecting biased ancestor/descendant relationships, while permuting between trees is more appropriate for detecting associations among traits (**Fig 3B**).

**Fig 3:**
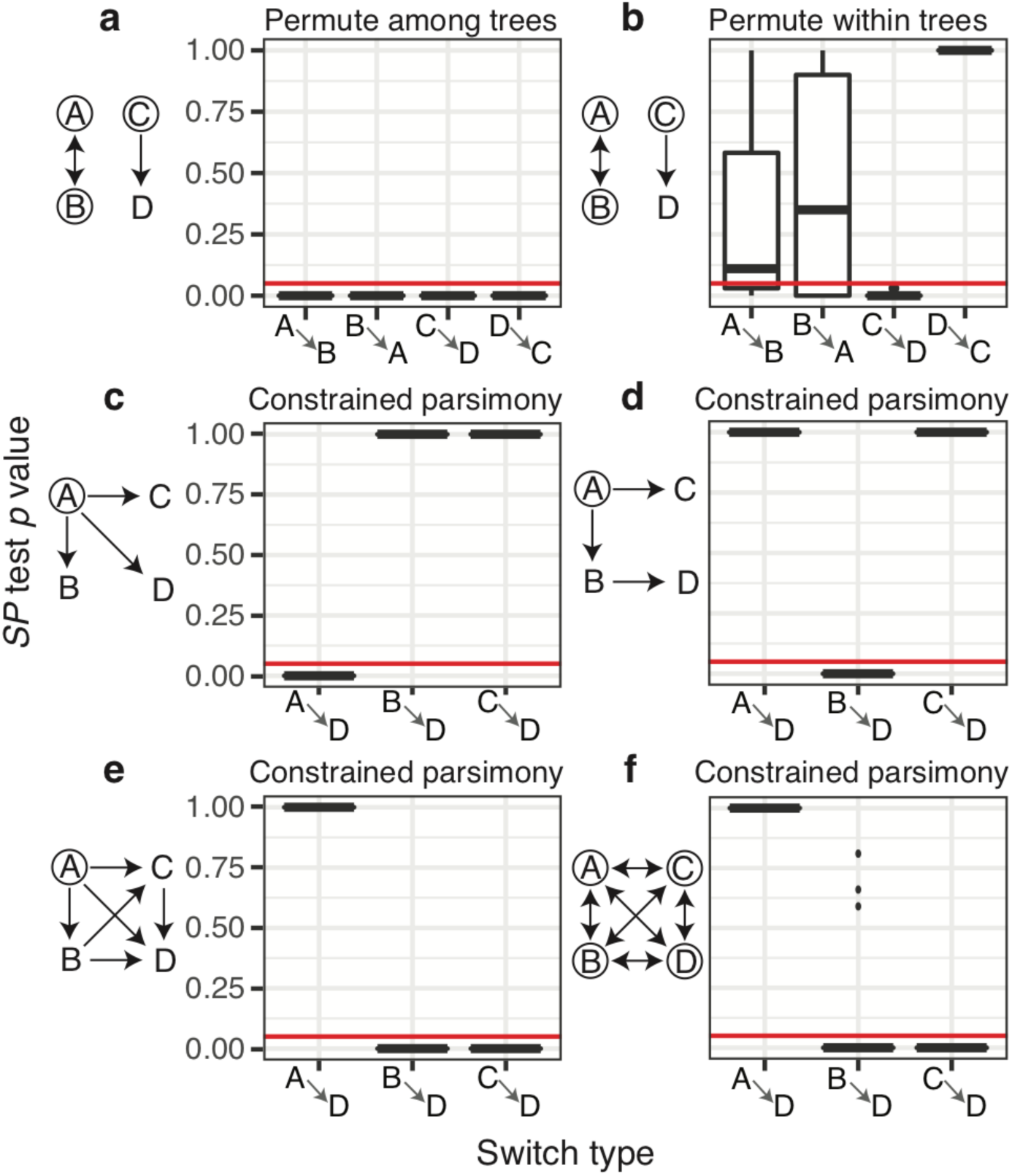
Distribution of *SP* test *p* values from four state simulation analyses under multiple modes of evolution diagrammed to the left of each plot. Twenty repetitions were performed in each scenario. In simulations, possible starting states are circled and possible state changes are shown with arrows. All allowed state changes occurred at the same relative rate and the total rate of state changing (*r*) was 10 changes/mutation/site (see **Fig 2**). **(a)** Permuting trait values among trees reveals low *p* values for all state changes between *A* and *B*, and *C* and *D*. **(b)** Permuting within each tree reveals low *p* values from *C* to *D*, but not between *A* and *B*. Both **a** and **b** imposed no constraints on the types of state changes allowed in the maximum parsimony algorithm. **(c)** Direct switching simulations result in low *p* values from *A* to *D*, but not from other states to *D*. **(d)** Sequential switching simulations result in low *p* values from *B* to *D* but not from other states to *D*. **(e)** Irreversible switching simulations result in low *p* values from *B* and *C* to *D*, but not from *A*. **(f)** Unconstrained switching simulations also result in low *p* values from *B* and *C* to *D*, but not from *A*. The strange results of **e** and **f** are likely artefacts of the constrained parsimony algorithm, which forbids reverse alphabetical state changes (e.g. *D* to *C*), used to count state changes in simulations **c-d**.

### Differentiating constrained modes of state change

In some instances, there are known constraints to the direction that state changes can occur, such as in isotype switching. Isotype-determining constant regions in humans are ordered as IgM/IgD, IgG3, IgG1, IgA1, IgG2, IgG4, IgE, IgA2. Human B cells begin with IgM/IgD, and because the mechanism of class switching is irreversible, these events can only occur sequentially in the order specified. For instance, IgA1 can switch to IgG4, but not to IgM or IgG1. This constraint may be naturally incorporated into the Sankoff parsimony algorithm [22] by making impossible isotype switches have an arbitrarily high weight. A frequent focus of isotype switching analysis is whether a particular isotype (i.e. IgE) arises from direct switching from IgM or from sequential switching from an intermediate isotype [30,31]. These types of hypotheses could be investigated using the *SP* test.

To determine if the *SP* test can usefully distinguish between types of constrained relationships among trait values, we simulated datasets to represent possible isotype switching patterns. As above, datasets contained four trait values: *A, B, C*, and *D* under different modes of evolution. Because questions often focus on the origin of a particular isotype [30] we only counted state changes leading to *D* when calculating *SP* statistics. Further, because state changes can only occur in a particular direction, we permute trait values among trees in these tests to increase power. While we previously showed that permuting among trees confuses biased association with biased ancestry (**Fig 3A**), switching between these states can only occur in one direction. Because of this, association between two states implies a direction of switching and among tree permutation is justifiable. We first simulate direct switching in which trees always had state *A* at the root and only state changes from *A* to the other states were allowed (**Fig 3C**). We expected these simulations to show a significantly high *SP* statistic only from *A* to *D*. Confirming this expectation, all 20 of these simulations had a high *SP* statistic from *A* to *D* (**δ** > 0, *p* < 0.05; **Fig 3C**). We next simulated sequential switching, where arriving at state *D* requires transitioning through *B*. All trees began in *A* and state changes were allowed from *A* to *B* and *C*, but *D* arose only from *B*. We expected these simulations to show a significantly high *SP* statistic only from *B* to *D*. All 20 of these simulations showed a significantly high *SP* statistic from *B* to *D* (**δ** > 0, *p* < 0.05; **Fig 3D**). These results demonstrate that the *SP* test using constrained parsimony can discriminate between simple hypotheses of isotype switch patterns, such as direct versus sequential switching.

We next investigated whether the *SP* test can distinguish between more complex types of constrained switching. We simulated irreversible isotype switching in which trees begin with state *A*, and only state changes moving alphabetically (*A* to *D*) were allowed. Naively, we may expect these simulations should show similar *SP* test results from *A, B*, and *C* to *D*. However, all 20 of these simulation repetitions showed a significantly high *SP* statistic to *D* from *B* and *C*, but not from *A* (**Fig 3E**). As a control, we simulated unconstrained switching in which trees begin randomly at any state and may change between all states. Using a constrained parsimony model, these simulations showed the same significantly high *SP* to *D* from *B* and *C*, but not from *A* (**Fig 3F**), indicating that this pattern is possibly an artifact of the constrained parsimony model. These results demonstrate that, while the *SP* test outlined here can distinguish between simple types of constrained state change, its relationship to more complex modes of constrained state change such as irreversible evolution are difficult to predict, and should be interpreted cautiously.

### Differentiation of B cell subtypes during HIV infection

Over the course of the immune response, B cells differentiate into multiple cellular subsets with distinct properties. Recent studies have focused on the role of T-bet, a transcription factor usually associated with differentiation of T cells, in shaping B cell responses during infection. For example, [6] used data from three HIV+ patients to demonstrate that CD19^hi^ memory B cells (CD19^hi^ MBCs, a surrogate for T-bet+ B cells) represented earlier states in the affinity maturation process than germinal center B cells (GCBCs), and to define the relationships among other B cell subtypes including CD19^lo^ MBCs and un-switched MBCs. More specifically, [6] used the *SC* test with trait values permuted among trees. However, the simulation analyses performed here demonstrated the *SC* test is highly conservative, and that permuting among trees may only detect unstructured association among trait values (**Fig 3**). It is therefore not clear whether the relationship from CD19^hi^ MBCs to GCBCs observed in [6] was driven by biased ancestor/descendant relationships among these cell types within trees. Our results above suggests that the *SP* test using within tree permutation would be a more appropriate test of this relationship.

We characterized the relationships among B cell subsets with the *SP* test using within tree permutation for each of the three subjects. These analyses showed a significantly high *SP* statistic from CD19^hi^ MBCs to GCBCs and to CD19^lo^ MBCs in all three subjects (**δ** > 0, *p* < 0.025; **Fig 4B-D**). These analyses confirm the conclusions in [6] that CD19^hi^ MBCs are significantly closer, cladistically, to the predicted germline sequence than GCBC sequences.

**Fig 4:**
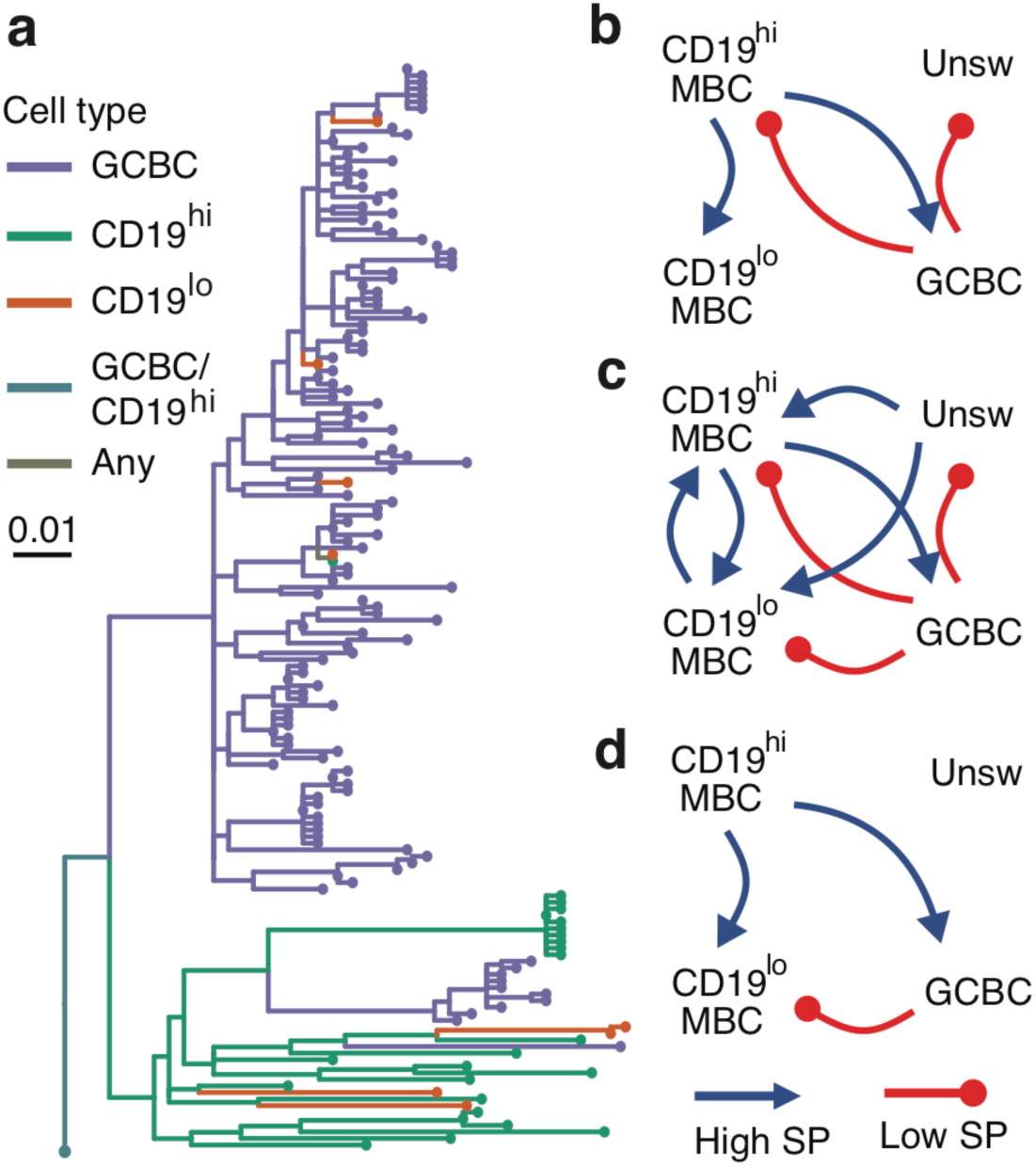
Analysis of B cell subtypes in three HIV+ subjects. **(a)** Example tree visualized using ggtree [37,38] showing observed relationship between CD19^hi^ MBCs and GCBCs. Ambiguous node states (“CD19^hi^/GCBC” and “Any”) are also shown. **(b-d)** Direction of significant *SP* test **δ** values for subjects 1 **(b)**, 2 **(c)**, and 3 **(d)**. Arrows within each diagram show the direction of significantly high (blue) or significantly low (red) *SP* statistics between CD19^hi^ MBCs, CD19^lo^ MBCs, unswitched MBCs (Unsw), and GCBCs in each subject.

Naively, one may interpret this as evidence that GCBC cells derive from CD19^hi^ MBCs. However, because GCBCs are expected to have far higher mutation rates than MBCs, the observed patterns are also consistent with early production of CD19^hi^ MBC from GCBCs, followed by a near cessation of mutations in CD19^hi^ MBCs. This is consistent with the conclusions of [6] that CD19^hi^ MBCs represent earlier stages in the GC reaction, rather than the direction of differentiation. Overall, these *SP* tests confirm that the previously observed relationships from CD19^hi^ MBCs and CD19^lo^ MBCs to GCBCs are driven by biased ancestor/descendant relationships within trees rather than simply association in the same trees, as may have been the case from the previously used *SC* tests with among tree permutations [6].

### Sequential isotype switching to IgE and IgG4

Antibody isotypes are a major determinant of function. Of principle interest is characterizing whether IgE antibodies, the primary antibody isotype associated with allergic response, arise directly from IgM switching, or through sequential switching from another downstream isotype [30,31]. Previous studies have shown evidence that IgE in mice and human adults arises from sequential switching primarily from IgG [30,31], though a recent study in 27 humans in the first three years of life found evidence of a greater association between IgA1 and IgE in children with food allergy and eczema [28]. Specifically, [28] showed a higher number of shared clones between IgE and IgA1 than between IgE and other isotypes in these subjects. A phylogenetic test of this relationship would confirm that IgE and IgA1 sequences show a direct ancestor-descendant relationship within these B cell trees rather than just being part of the same clone.

We applied our discrete trait framework to determine the origins of IgE in a single subject (id = 2442) from [28]. This subject was selected due to reported history eczema, food allergy, and B cell clones containing IgE and other isotypes [28]. Using an *SP* test in which only state changes leading to IgE were considered and trait values were permuted among trees, we found a significantly high *SP* statistic from IgA1 to IgE (**Fig 5B**). No other isotype showed a significantly high *SP* statistic to IgE. These results favor IgE arising from sequential switching through IgA1 over direct switching from IgM in this subject. Performing a similar test using only state changes leading to IgG4 revealed a significantly high *SP* statistic from IgG1 and IgG2 to IgG4 (**Fig 5E**). This pattern is similar to irreversible switching within the IgG family (**Fig 3E**). As shown in simulation analyses, this test is not suited to infer relative rates of switching from different isotypes if all kinds of switches are considered. However, these results are most consistent with origin of IgG4 through sequential switching with other IgG isotypes rather than direct switching from IgM or sequential switching from IgA1. Overall, these results are consistent with the conclusions of [28] that IgE arose preferentially through switching from IgA1 in this subject. Our results further suggest IgG4 arose preferentially via sequential switching from other IgG subtypes in this subject.

**Fig 5:**
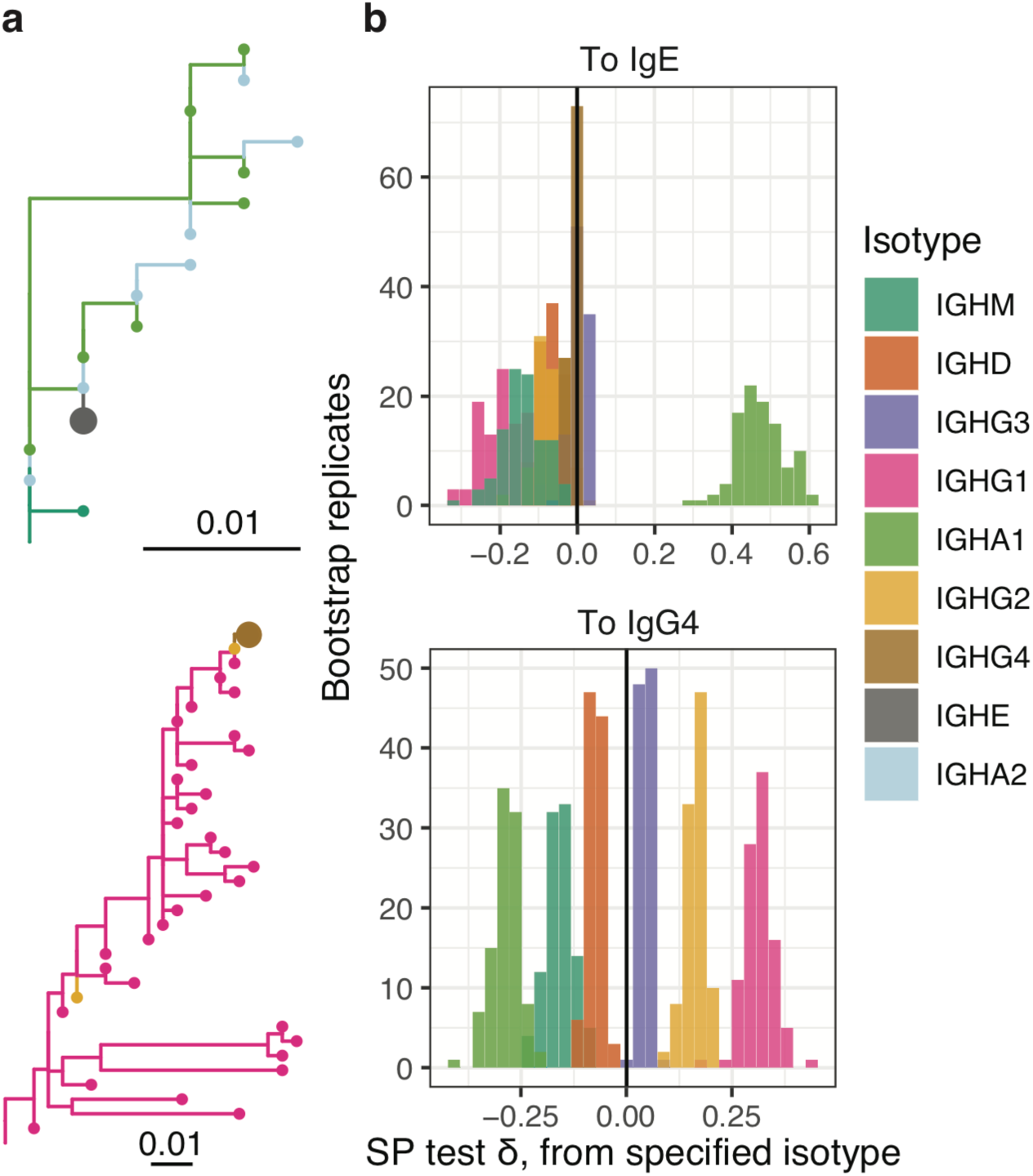
Analysis of antibody isotypes from a single subject. **(a)** Example trees visualized using ggtree [37,38] showing observed relationships between cells expressing BCRs with IgA1 and IgE, and IgG1 and IgG4 isotypes. IgE and IgG4 are indicated on each tree using larger tip circles. **(b)** Distribution of *SP* test δ values to IgE from each of the other isotypes (different colors). **(c)** Distribution of *SP* test δ values to IgG4 from each of the other isotypes (different colors). State changes in **b** and **c** were calculated using constrained parsimony which forbids state changes that violate the geometry of the Ig heavy chain locus, and *SP* tests were performed using permutation among trees.

## Discussion

Phylogenetic techniques have the potential to reveal important information about B cell migration, class switching, and cellular differentiation. While great strides have been made using phylogenetic models to study evolution and trait change generally, there are significant challenges to translating these approach to B cells. As a step in this direction, hypotheses about the ancestor/descendant relationships of B cell trait values may be usefully investigated using heuristic approaches that are robust to uncertainties in branch length estimation. Here, we introduce three maximum parsimony-based summary statistics to characterize the distribution of trait values along phylogenetic trees. Significance of all of these statistics is tested by comparing to statistics calculated on trees with permuted data. We demonstrate the efficacy of these tests using simulations, and show that the *SP* test is the most useful for characterizing ancestor/descendant relationships among trait values. We further demonstrate how these statistics can test hypotheses about empirical B cell datasets by characterizing the relationship between T-Bet+ memory B cells and germinal center B cells in three HIV+ patients, and the class switching origins of IgE and IgG4 in a human subject over the first three years of life.

Simulations demonstrate that the *SP* test was uniquely able to determine the direction of biased origination and state change among the approaches investigated. In simple simulations containing two states (*A* and *B*) a significantly high *SP* statistic from *A* to *B* was associated with origination in *A* and biased state change from *A*. This signal decreased as the overall rate of switching increased. In more complex scenarios, the *SP* test was able to differentiate between traits that were generated through biased state change in a particular direction versus traits that were simply associated with each other. The *SP* test was also able to distinguish between simple modes of constrained evolution such as direct and sequential switching. These results indicate the *SP* test may have broad utility in characterizing ancestor/descendant relationships among B cell discrete traits.

We next used two datasets to demonstrate that the *SP* test could be used to derive meaningful biological conclusions. In the first, we confirm that T-Bet+ memory B cells tend to be the predicted immediate ancestors of GC B cells within lineages trees obtained from three HIV+ subjects. Though this relationship may primarily be due to differences in mutation rate over time between memory and GC B cell subsets, this does confirm prior findings and demonstrates that the T-Bet+ memory B cell subset represents an earlier state in the affinity maturation process, possibly contributing to an impaired immune response to HIV [6]. We next characterized the isotype switching patterns of sequences obtained from a human over the first three years of life [28]. In this analysis, we found evidence of sequential switching from IgA1 to IgE, as well as evidence of sequential switching from IgG subtypes to IgG4. Sequential switching from IgA1 to IgE is consistent with [28] but not other analyses performed on data taken from adults, which favor sequential switching from IgG [30]. This possibly reflects differences in isotype switching patterns between adults and children. Overall these results demonstrate that the discrete trait analysis framework developed here can be used to test important hypotheses about B cell differentiation and class switching.

There are a number of limitations with these methods. First, tree topologies were estimated using maximum parsimony. While maximum parsimony is not a statistically consistent estimator of tree topology and is known to give inaccurate predictions over long branch lengths [32], it has been shown to be an accurate estimator of tree topology in certain B cell applications [33], and is widely used in B cell phylogenetic analysis [5,17]. In any case, the statistics presented here are not limited to tree topologies inferred through maximum parsimony. The three statistics proposed here (**Eq. 1-3**) are also based on maximum parsimony, and may have similar inaccuracies over long branch lengths. Further, the statistical tests assume that the process of state change is independent of the tree shape, when the two may be coupled e.g. [34]. This assumption of independence is commonly made in discrete trait analysis e.g. [8] to enable computational tractability, and because the actual link between tree shape and state change is unknown or cannot be modelled. A significantly high *SP* statistic could be potentially caused by factors other than biased state change. For instance, because tree branch lengths represent genetic distance rather than time, it is possible that cell types with low mutation rates over time will spuriously appear ancestral to those with high mutation rates. This effect likely underlies our analysis of B cell subtypes in HIV. Finally, it is possible that SHM is actually occurring at another, un-sampled site which is seeding the sites that were sampled. Overall, it is important to carefully consider alternative explanations when trying to determine the biological basis for a significantly high *SP* statistic.

An important limitation of the *SP* test is that it, like many other phylogeographic approaches e.g. [29] is affected by biased data sampling. This may arise due to experimental factors that are difficult to control. For instance, under-sampling a trait value may cause a spurious, significantly high *SP* from that trait value. Previous analyses of viral migration have dealt with potential sampling bias by performing tests across multiple down-sampling repetitions [9]. In practice, it can be difficult to know if B cells with certain trait values have been sampled proportionally to their relative population sizes. However, if a type of B cell is known to be under-sampled in a particular experiment, and is predicted to be the descendant of another B cell type, it can be argued that this relationship is unlikely to be due to biased sampling (**Fig 2E**). Alternatively, if multiple samples are tested it is possible that these samples will have a wide range of sequence proportions belonging to different traits. If these differences in sequence proportions are uncorrelated with *SP* test results, it could be argued that observed results are unlikely to be due to consistent under-sampling of B cells with a particular trait value.

Our simulation analyses revealed that that the *SP* test is difficult to interpret when considering complex constrained models such as irreversible isotype switching (**Fig 3C-E**). To recreate isotype switching, we performed four state simulations in which only state changes proceeding in the direction of state *A, B, C*, and *D* were allowed. Unexpectedly, these simulations tended to show a significantly low *SP* statistic from *A* to *D*, but a significantly high *SP* statistic for *B* and *C* to *D* (**Fig 3E**). This biased trend is likely driven by the fact, due to constraints in the direction of state change, randomized trees tend to have more switches from *A* than expected based on the relative frequency of *A*. This produces a significantly high *SP* for switches from *A* to *D*. An alternative may be to use the *SP* statistic (**Eq. 3**) without comparing to a null distribution, which is equivalent to comparing the relative frequency of each type of switch observed. However, the observed switch frequency (*SP* statistic) is not proportional to the true relative rate of state change in general. For instance, in the two state Markov model simulations presented here (**Fig 2**), the *SP* statistic alone is both positively and negatively related to the true relative rate of state change, depending on other parameters (**Supplemental File S5**). Comparing *SP* statistics to those obtained from randomized trees (i.e. the *SP* test) usefully corrects this relationship in unconstrained models (**Fig 2**), but not always in constrained ones (**Fig 3C-E**). Ultimately, isotype switching is a complex, constrained process, and our analyses suggest the relative rates of isotype switching inferred from B cell trees should be interpreted cautiously. We suspect a general method for accurately estimating these rates will require a model-based approach, such as a non-reversible Markov model.

Future methods to differentiate migration, differentiation, and isotype switching patterns in B cells might improve upon the approach developed here by explicitly modeling these processes along a phylogeny, incorporating branch length information, and better accounting for uncertainty in tree topology. The heuristic approach introduced here crucially does not use branch lengths to help predict internal node states of the tree. Ignoring this source of information likely lowers power, but is possibly advantageous because the relationship between mutation rate and time is not currently well understood, and likely varies by cell type. While the approach developed here uses phylogenetic bootstrap replicates to account for uncertainty in tree topology [24], this may also be done using a posterior distribution of topologies generated by MCMC sampling. This was recently done for naïve sequence inference in individual B cell lineages [35]. Phylogenetic bootstrapping has less desirable statistical properties than posterior distributions, but is a widely used means of assessing reproducibility of tree topology and is more computationally tractable for large datasets. Overall, though there is potential for improvement, the approach introduced here effectively deals with important challenges such as incorporating information across trees, accounting for uncertainty in tree topology, and scaling efficiently when analyzing large datasets.

A phylogenetic discrete trait analysis framework fills an important gap in B cell sequence analysis. The proposed framework provides a principled, flexible, and scalable approach for characterizing migration, class switching, and differentiation in a wide array of contexts. This differs from other phylogenetic tools we developed recently, which used model-based approaches for characterizing somatic hypermutation and clonal selection [4,36]. The methods developed in this paper are available in the R package *dowser*, available at https://bitbucket.org/kleinstein/dowser as part of the Immcantation suite (http://immcantation.org). Scripts for performing analyses in this manuscript are available at: https://bitbucket.org/kleinstein/projects.

## Acknowledgements

K.B.H was supported through a PhRMA Foundation post-doctoral fellowship in informatics. This work was funded in part by the European Research Council under the European Union’s Seventh Framework Programme (FP7/2007-2013)/European Research Council grant agreement number 614725-PATHPHYLODYN. This work was funded in part by National Institutes of Health, National Institute of Allergy and Infectious Diseases grant R01 AI104739.

