## Supplemental Information for "Phylogenetic analysis of migration, differentiation, and class switching in B cells"

**Supplemental File 1:** Maximum ambiguity resolution of polytomies

B cell lineage trees frequently have clusters of nodes separated by zero-length branches called polytomies. Polytomies represent a high degree of uncertainty in ancestor-descendant relationships within phylogenetic trees (**Fig S1a**) and can be resolved into a multiple distinct, equally supported sets of bifurcating nodes (**Fig S1b**). We took several steps to ensure that polytomies within each tree were resolved in a manner that appropriately represents this uncertainty. First, all polytomies were re-ordered using nearest-neighbor interchange (NNI) moves to minimize the number of label switches along the tree beginning with the germline sequence and moving down (**Fig S1c**). Then, each polytomy was arranged to a maximum ambiguity configuration, which has the largest number of distinct maximally parsimonious node switching histories (**Fig S1d**). To accomplish this, we used the Sankoff parsimony algorithm to label the state at each internal node of the tree. For each polytomy, we then removed all zero-length branches connecting the ancestral node of the polytomy (*A*; **Fig S1d** node 1) with the descendant nodes of the polytomy (*D*). We then joined all *D* nodes with the same state into separate subtrees, and attached the upper-most nodes of these new single-type subtrees (*T*; **Fig S1d** nodes 3, 4, and 5) to node *A* in a balanced manner. To represent switch types across ambiguous internal node sets, maximum parsimony trajectories were randomly chosen in the backtrace step of the Sankoff algorithm. This process moves down the tree beginning at the root node, and is performed 100 times for each tree. The mean value of each switch type across these repetitions is reported. However, for polytomies with more than two states, there are multiple distinct ways the nodes *T* could be joined to node *A*. To account for this, for each repetition of the sampling procedure, the nodes *T* within each resolved polytomy are randomly swapped using an NNI move. Further, switches were not counted between polytomies and their descendant nodes *D*; rather, the state of each descendant node was randomly chosen from all states contained in the polytomy which could be that node's ancestor with equal parsimony (**Fig S1d** node 6). These procedures ensured that the minimum number of label switches occurred within each polytomy, and that all distinct sets of label switches along the polytomy were appropriately represented. This method has been implemented in the B cell phylogenetic software package IgPhyML v1.1.1 (bitbucket.org/kleinstein/igphyml).

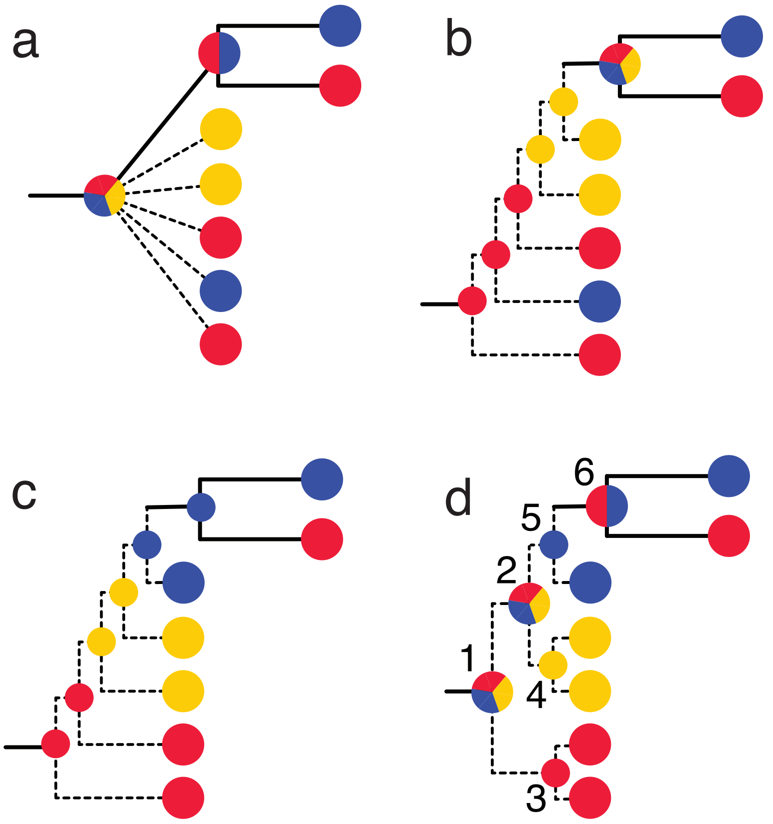


**Fig S1**: Representations of a polytomy. Trait values are represented by red, blue, and yellow colored nodes. Dashed lines represent zero-length branches (no mutations), while solid lines represent nonzero-length branches. Node colors show maximum parsimony predicted trait values. Multi-colored nodes represent multiple equally parsimonious states. **a)** A polytomy represented using a multifurcating node. **b)** Random resolution of the polytomy into a binary tree gives more switches than needed (4), and places red at the root. **c)** Maximum parsimony resolution reduces the total number of switches (3) but, this configuration places the order of switching along the tree as red, yellow, blue, red, when in reality the order of switching cannot be determined. This leads to a switch count of red-yellow=1, yellow-blue=1, blue-red=1. **d)** Maximum parsimony and maximum ambiguity configuration appropriately represents possible switching scenarios under maximum parsimony. The root node (node 1) can be either color with equal parsimony score (3). Switching histories are sampled, starting at node 1 and moving down the tree. Switches are not counted across nonzero branches leading from the polytomy (e.g. between node 5 and node 6). Instead, the state at node 6 is randomly chosen among maximally parsimonious states represented by the polytomy (i.e. red and blue). Further, nodes 3, 4, and 5 are swapped each sampling repetition to ensure each state as equal probability of being at the root node. After multiple sample repetitions, the average number of each type of switch is recorded. The 3 switches that occur are distributed accordingly: red-blue=0.83 switches, blue-red=0.83, red-yellow=0.33, yellow-red=0.33, yellow-blue=0.33, blue-yellow=0.33.

**Supplemental File 2:** Processing of empirical datasets

Our isotype analysis dataset was aimed at understanding isotype switching patterns in human infants, and consists of BCR mRNA sequences obtained from peripheral blood samples taken from a human child each year from age 1 to 3 years old [1]. Preprocessing was performed with pRESTO v0.5.13 [2]. Quality control was performed by first removing all sequences with a Phred quality score < 20, length < 300bp, or any missing ("N") nucleotides. The 3' and 5' ends of each read were matched to forward and constant region primers with a maximum error rate of 0.1. The region adjacent to the constant region primer was exactly matched to sub-isotype specific internal constant region sequences obtained from [1]. Only sequences with the same isotype predicted by their constant region primer and internal constant region sequence were retained. Identical reads were collapsed and identical sequences observed only once were discarded. V(D)J assignment was performed using IgBLAST v 1.13 [3] against the IMGT human germline reference database (IMGT/GENE-DB v3.1.24; retrieved August 3rd, 2019; [4]). Putatively non-productively rearranged sequences were removed. To infer clonal clusters, sequences were first partitioned based on common IGHV and IGHJ annotation, and junction region length. Within these groups, sequences differing from one another by a normalized Hamming distance of 0.1 within the junction region were clustered into clones using single linkage hierarchical clustering [5]. The V and J genes of unmutated germline ancestors for each clone were then constructed with D segment and N/P regions masked by "N" nucleotides. Clonal clustering and germline sequence reconstruction were performed with Change-O v0.4.6 [6]. Error resulting from repeated sequencing of the same molecule was reduced using a similar approach to [7]. Namely, sequences were removed if they differed by a Hamming distance of 1 from another sequence found 100 times more frequently, or if they differed by Hamming distance of 2 from another sequence found 1000 times more frequently, and so on following a frequency ratio cutoff of $10^{(\text{Hamming distance}+1)}$.

**Supplemental File 3:** Simulation analyses with *PS* test

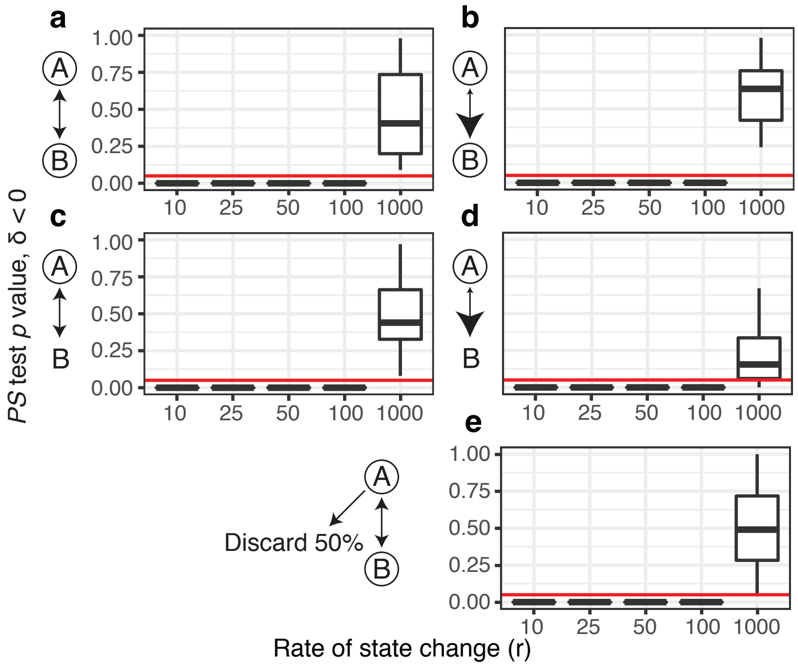


**Fig S3**: Distribution of *PS* test *p* values for the hypothesis that **δ** < 0 from two state simulation analyses. See **Fig 2** for analysis of the same data with the *SP* test. In these simulations, change between state *A* and *B* was determined by the probability of starting in *A* ($\pi_a$), relative rate of migrating from *A* to *B* ($r_{ab}$), and the average rate of state change ($r$). To the left of each plot, possible starting states are circled, relative rates are shown by arrowhead size. **(a)** $\pi_a = 0.5$, $r_{ab} = 1$, fully unbiased state change. **(b)** $\pi_a = 1$, $r_{ab} = 1$. **(c)** $\pi_a = 0.5$, $r_{ab} = 10$. **(d)** $\pi_a = 1$, and $r_{ab} = 10$. **(e)** $\pi_A = 1$, $r_{ab} = 1$, 50% of *A* sequences are discarded. Red lines show the cutoff of $p$ value = 0.05.

**Supplemental File 4:** Simulation analyses with *SC* test

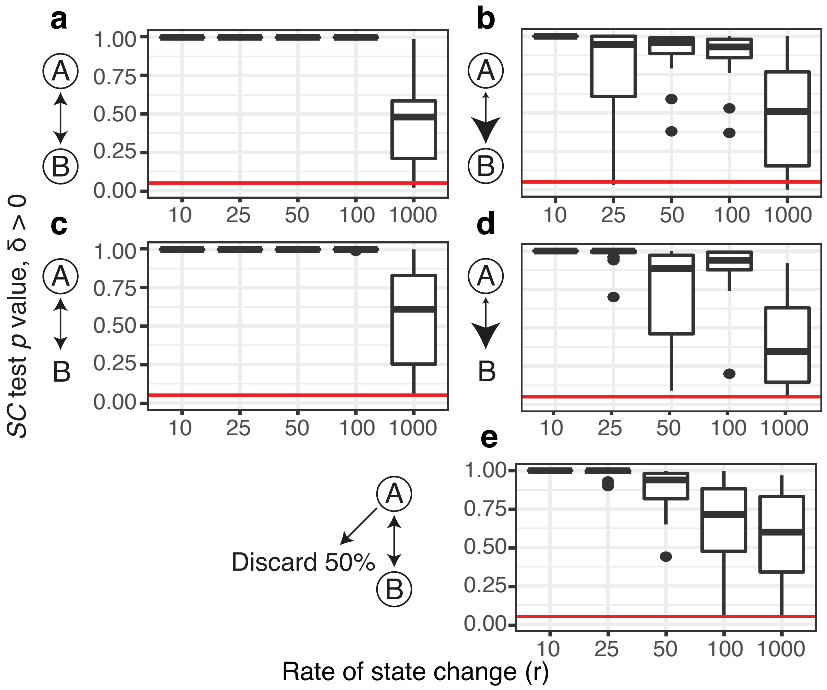


**Fig S4**: Distribution of *SC* test *p* values for the hypothesis that **δ** > 0 from two state simulation analyses. See **Fig 2** for analysis of the same data with the *SP* test. In these simulations, change between state *A* and *B* was determined by the probability of starting in *A* ($\pi_a$), relative rate of migrating from *A* to *B* ($r_{ab}$), and the average rate of state change (*r*). To the left of each plot, possible starting states are circled, relative rates are shown by arrowhead size. **(a)** $\pi_a = 0.5$, $r_{ab} = 1$, fully unbiased state change. **(b)** $\pi_a = 1$, $r_{ab} = 1$. **(c)** $\pi_a = 0.5$, $r_{ab} = 10$. **(d)** $\pi_a = 1$, and $r_{ab} = 10$. **(e)** $\pi_A = 1$, $r_{ab} = 1$, 50% of *A* sequences are discarded. Red lines show the cutoff of *p* value = 0.05.

**Supplemental File 5:** *SP* statistics in two state simulation analyses

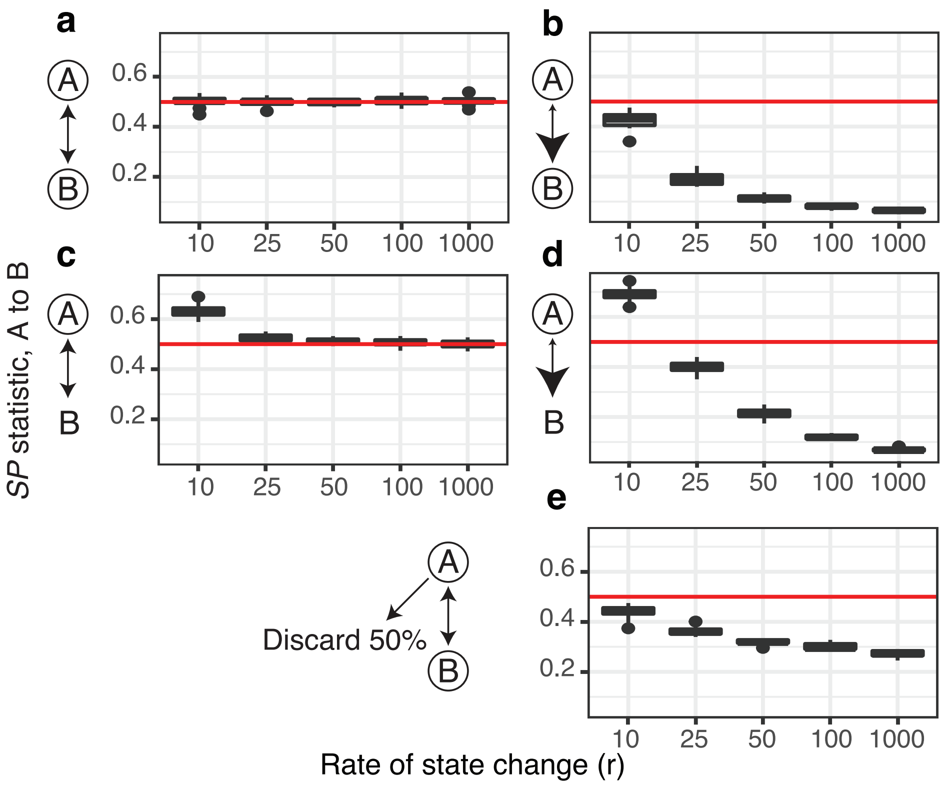


**Fig S5**: Distribution of raw *SP* statistics from *A* to *B* in two state simulations. The red line is at 0.5, showing equal switch frequency. See **Fig 2** for analysis of the same data with the full *SP* test. In these simulations, change between state *A* and *B* was determined by the probability of starting in *A* ($\pi_a$), relative rate of migrating from *A* to *B* ($r_{ab}$), and the average rate of state change ($r$). To the left of each plot, possible starting states are circled, relative rates are shown by arrowhead size. **(a)** $\pi_a = 0.5$, $r_{ab} = 1$, fully unbiased state change. **(b)** $\pi_a = 1$, $r_{ab} = 1$. **(c)** $\pi_a = 0.5$, $r_{ab} = 10$. **(d)** $\pi_a = 1$, and $r_{ab} = 10$. **(e)** $\pi_A = 1$, $r_{ab} = 1$, 50% of *A* sequences are discarded. Note that at low rates ($r = 10$), origination at *A* (**c** and **d**) shows *SP* statistic > 0.5. However, at higher rates ($r > 10$), increased rate of state change from *A* to *B* actually produces lower *SP* statistics. This effect increases as overall rate ($r$) increases. This indicates that the *SP* statistic itself is not a good estimator for the relative rate of state change, and is why we only interpret *p* values resulting from comparison of the *SP* statistic to a null distribution (i.e. the *SP* test).
